## Supplementary Figures for "INSERT-seq enables high resolution mapping of genomically integrated DNA using nanopore sequencing"

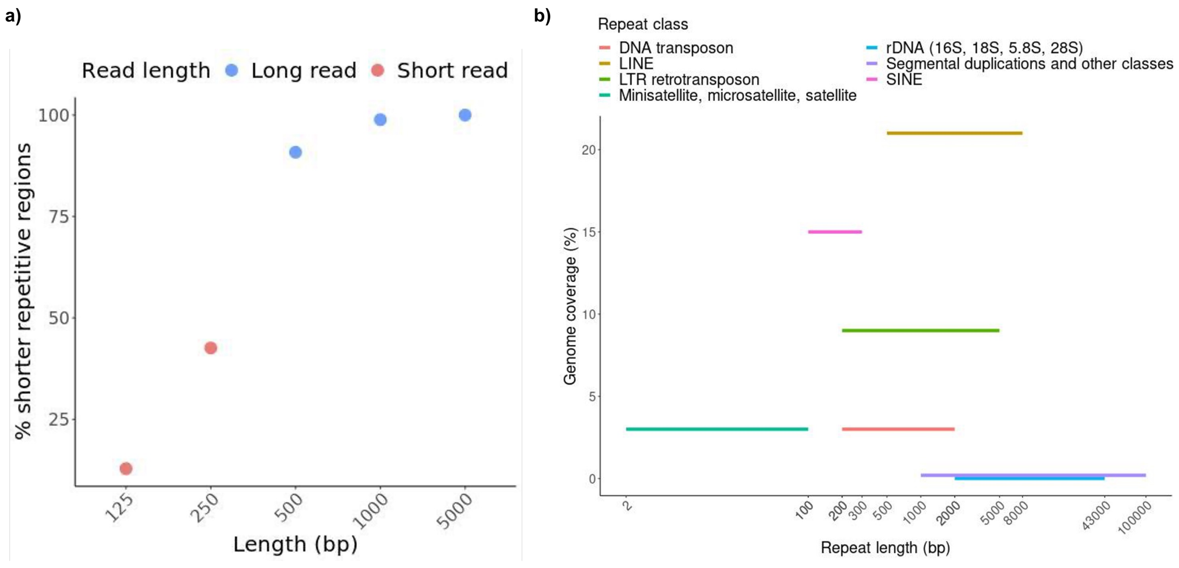


**Supplementary Figure 1 | a)** Percentage of Dfam database human genome repetitive regions shorter than 125, 250, 500, 1000 and 5000 base pairs. **b)** Representation of repetitive regions from Human genomes based on their length range in the x axis and the percentage of representation in the genome in the y axis.

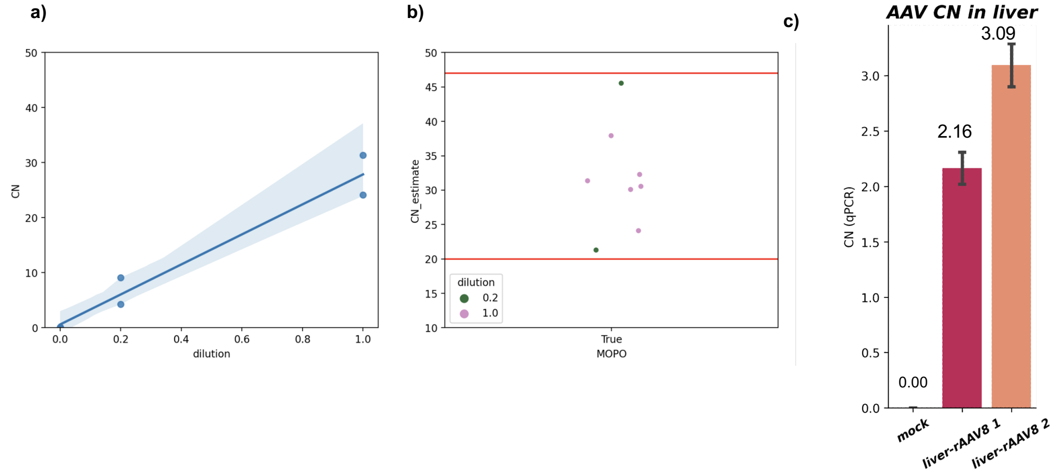


**Supplementary Figure 2 | CN determination a)** Determination of integrated LV copy number in mopo number. MOPO DNA and WT genomic DNA where mixed in different ratios to asses linearity of the ddPCR method . **b)** Diferent CN estimates of the same MOPO sample
**c)** Quantification of AAV DNA levels in mouse liver by qPCR. Numbers indicate two different animal replicates.

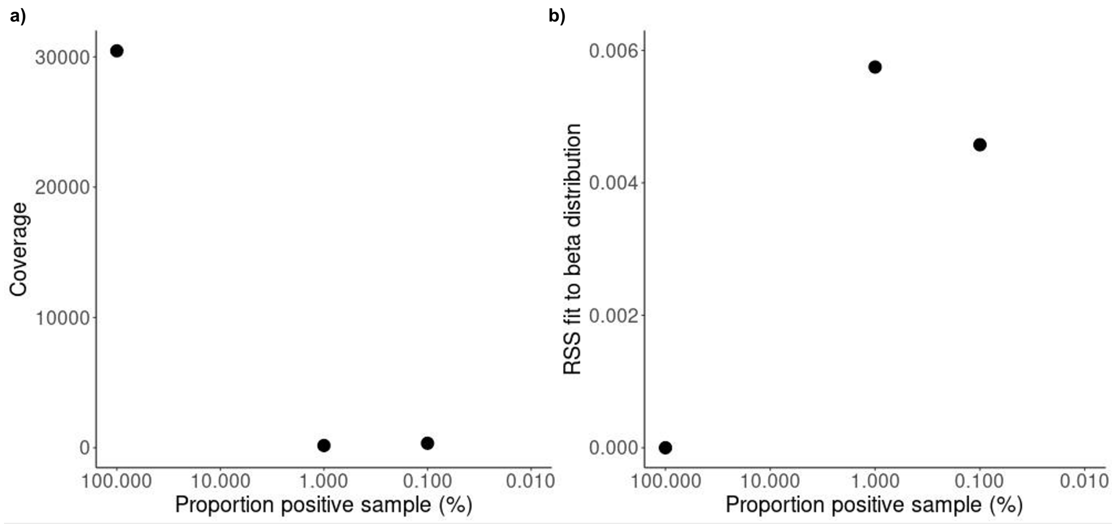


**Supplementary Figure 3 | Limit of Detection (LOD) calculation. a)** Insertion detection depending on the sample dilution. Shows the coverage of on-target insertions in a sample without dilution, diluted 1:100, 1:1000 and 1:10000. **b)** Shows the Residual Sum of Squares (RSS) from the fit of insertion coverage to a beta distribution from detected on-target insertions in a sample without dilution, diluted 1:100, 1:1000 and 1:10000.

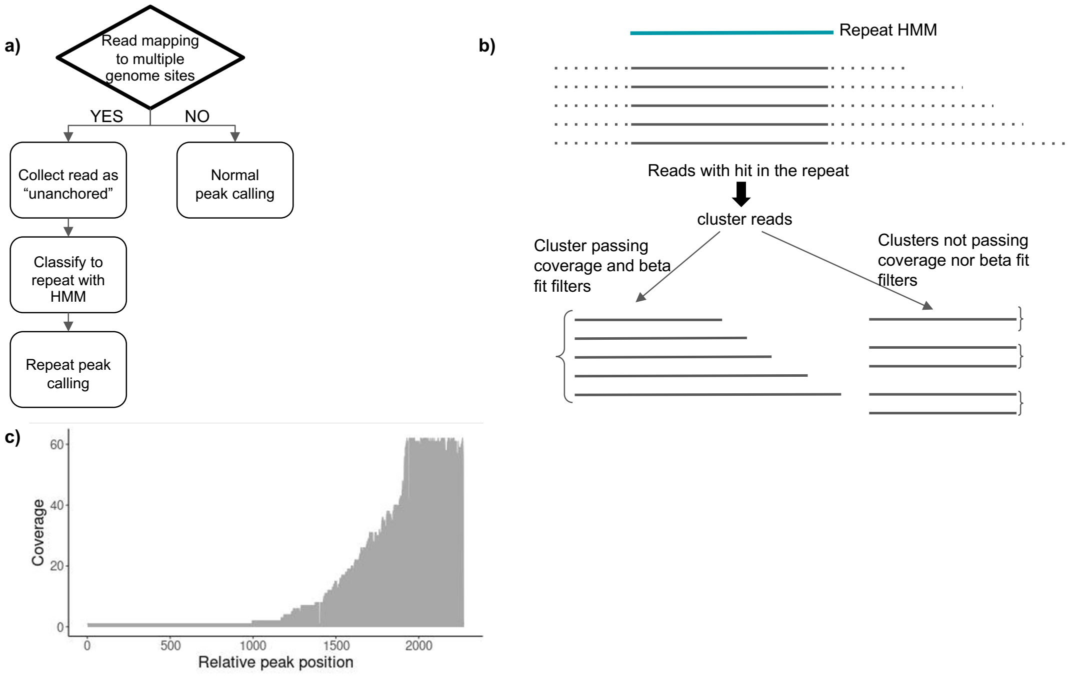


**Supplementary Figure 4 | Unanchored peak calling pipeline. a)** Schematic representation of the detection of insertions at repetitive regions. Reads that map to multiple genome sites are collected and mapped to the reference human repeats. A peak calling of the repeats is performed. **b)** Schematic representation of the peak calling of unanchored reads. Unanchored reads are classified to a repeat based on HMM hits, reads from each peak are clustered in order to determine if all the reads belong to the same peak, finally, a coverage filter and standard deviation

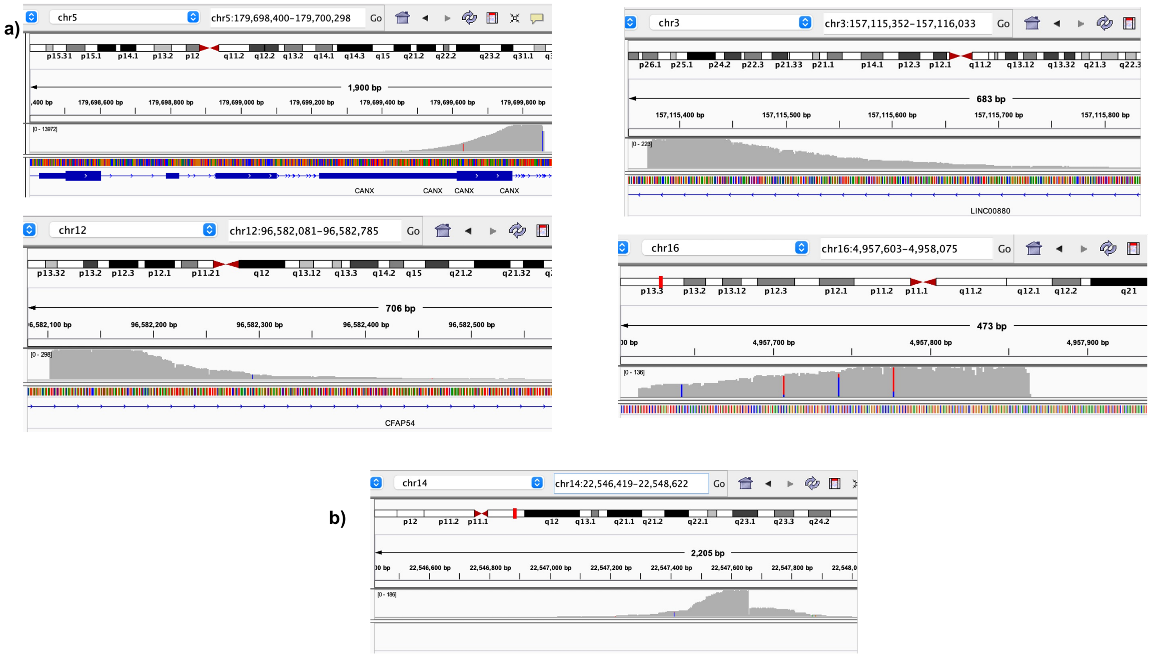


**Supplementary Figure 6 | Integration profile of Cas9-PiggyBac chimeric protein:** Coverage at 5 most represented integration sites at **a)** off-target insertions and **b)** on-target insertion. Off-targets in such abundance but occurring at the same location could be originated from clonal expansions.

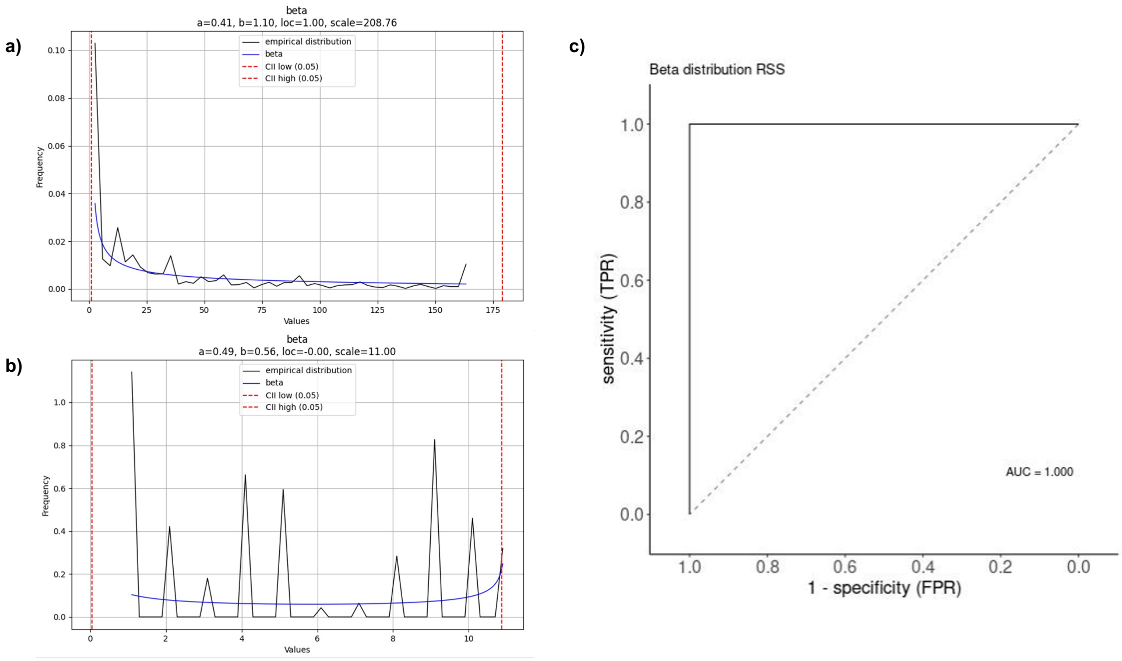


**Supplementary Figure 8 | Peak calling ROC curve of MOPO sample. a)** Representation of the distribution of coverage (black) of a true peak from the MOPO sample at chr15:99327982-99330675 fitted to a beta distribution (blue) with RSS=0.005045. **b)** Representation of the distribution of coverage (black) of a false peak from the MOPO sample at chr3:99327982-99330675 fitted to a beta distribution (blue) with RSS=2.831412. **c)** ROC curve of the Residual Sum of Squares (RSS) of peak coverage fit to a beta distribution. The selected threshold was 1.

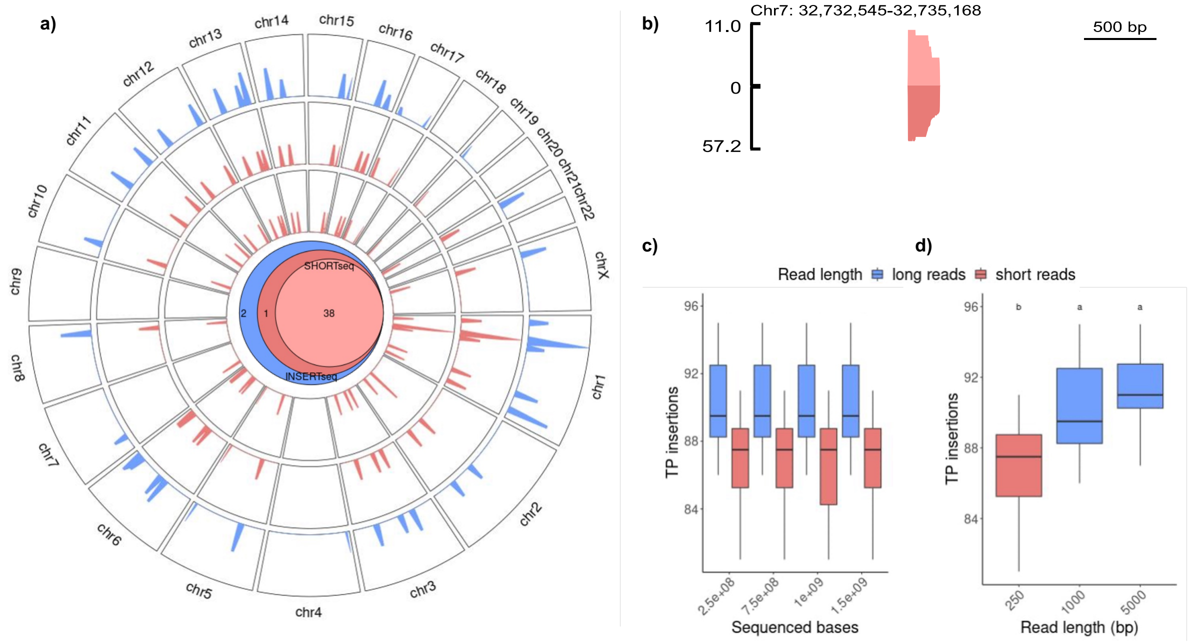


**Supplementary Figure 9. | Effect of increased number of reads. a)** Genome-wide map displaying the overlap between insertions detected in MOPO with either short read insertion detection (red) and long read insertion detection (blue). The Venn diagram summarises the overlap of common insertions with the two methods. A higher number of reads was used for a SHORTsesq analysis (dark red) where one new insertion is found respect to the experiment with a subset of reads. **b)** Coverage at insertion site in the mono-clonal poli-insertional (MOPO) analyzed cell line, with a subset of short reads (light red) and the complete run of short reads (dark red). The insertion shown was detected with a higher sequencing depth. **c)** Number of true positive (TP) insertions detected in a simulated dataset of 100 random insertions and 10 replicates, with reads of either 250bp and 1000bp and a total number of sequenced bases from 250Mb to 1.5Gb. **d)** Number of true positive (TP) insertions detected with read length of 250, 1000 and 5000 base pairs. Statistically significant differences (p<0'05) are annotated by groups with letters (a, b).

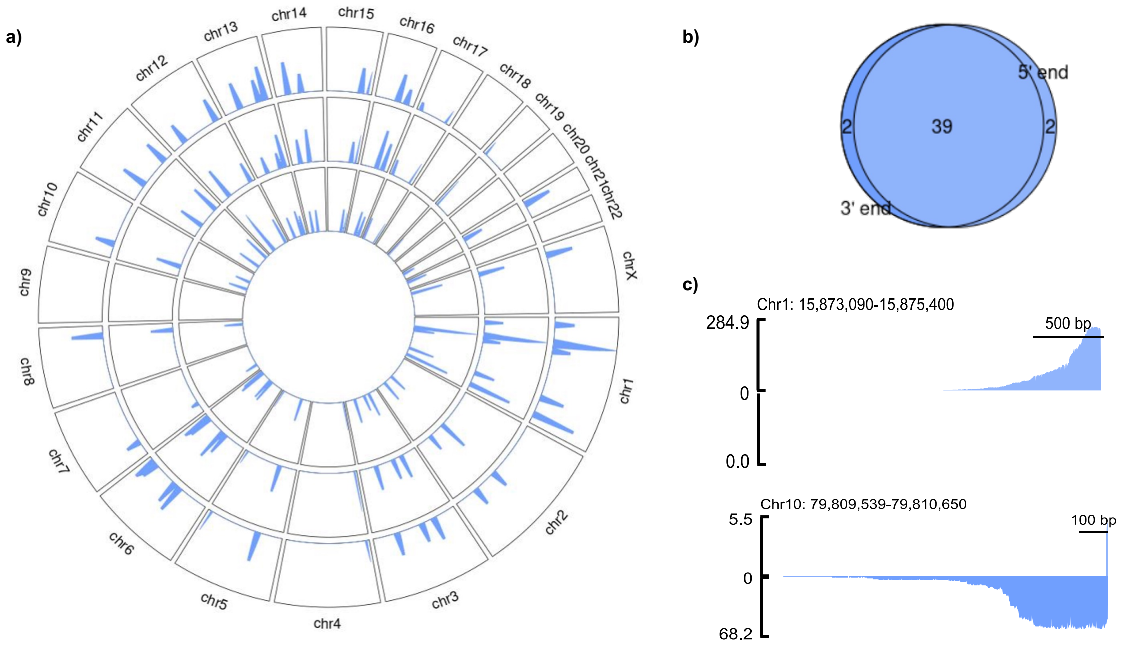


**Supplementary Figure 10. | Analysis of sample replicates. a)** Genome-wide map displaying the overlap between insertions detected in MOPO long read insertion detection. Insertion mapping was performed to both lentiviral ends 5’ (two different replicates) and 3’. **b)** The Venn diagram summarises the overlap of common insertions with the two methods. Two insertions are found with the 5’ end mapping and not with the 3’ end while 2 new insertions are found only with the 3’ end mapping. **c)** Coverage at two selected insertion sites in the mono-clonal poli-insertional (MOPO) analyzed cell line, with 5’ end mapping (light blue) and 3’ end mapping (dark blue). Two representative insertion sites detected with only one of the end mappings are shown.

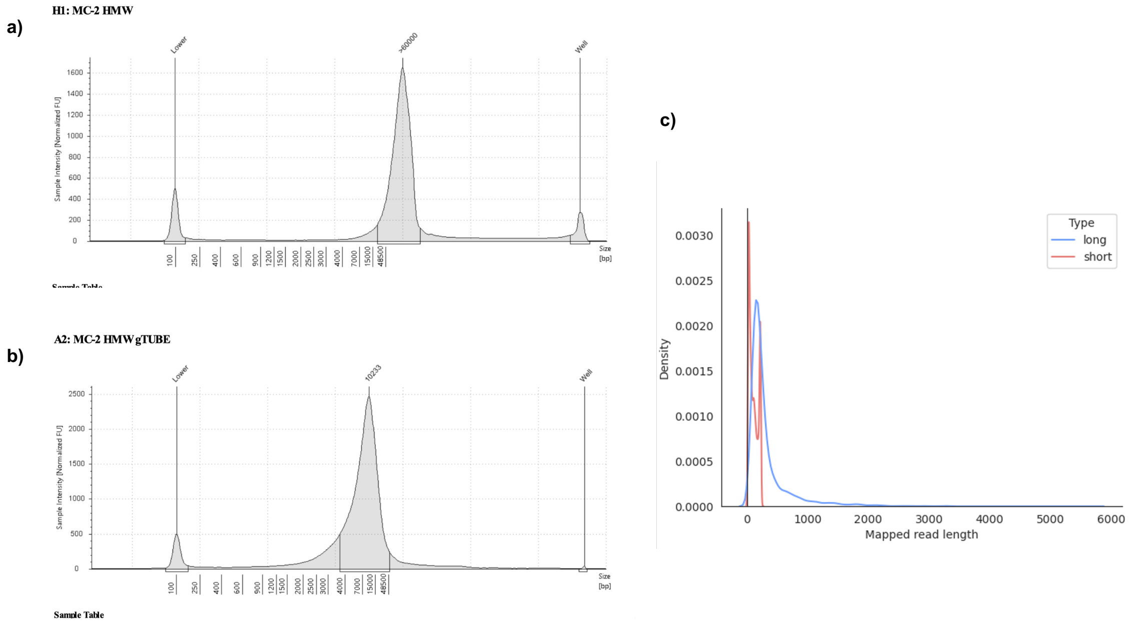


**Supplementary Figure 11. | Library prep impact on read length. a)** Bioanalyzer showing the distribution of fragment length of the MOPO sample before random fragmentation. Mean fragment length was higher than 60 kbp. **b)** Bioanalyzer showing the distribution of fragment length of the MOPO sample after random fragmentation. Mean fragment length was ~10 kbp **c)** Density distribution of mapped read length of the MOPO sample.
