## Supplementary Protocol for "INSERT-seq enables high resolution mapping of genomically integrated DNA using nanopore sequencing"

Benchling

UPDATED 8/6/2021 01:17 PM

by Marc Güell (created by dimi)

### Introduction

The described protocol allows retriving on/off targett patterns of a genomic perturbations of >20bp in size. Using STAT-PCR (Tsai. et al 2015) Nanopore sequencing allows for retrival of longer reads that facilitate mapping of the flanking regions of the insert by bypassing repetitive genome elements. This protocol uses ONT PCR Barcoding Kit (SQK-$PBK_{004}$)

### Materials

- DNA Extraction KIT
- gTUBE / sonicator
- KAPPA HYPERPREP KIT (or end repair and T4 lig NEB)
- ISA## ADAPTORS
- MISEQ comon ADAPTOR
- $P_{05}$ oligos
- PBK insert specific oligos
- PBK ONT KIT
- LongAmp Hot Start Taq 2X Master Mix
- Thermocycler
- MiniON sequencer
- lambda Exo from NEB

### Procedure

#### DNA extraction

1. Perform DNA extraction using either QUIAGEN COLUMN or CIRCULOMICS or column method.

#### Library preparation

2. Fragment DNA 35s at 12000 RPM, using 15µg of DNA, using gTube, and based on gTUBE protocol.
3. Set up End repair using 50µL (5µg) of gTUBED DNA following above table:

**End Repair**

| | A | B | /6 | for 16 | |
|---|---|---|---|---|---|
| 1 | Component | Volume | | | |
| 2 | H20 | 0.00 | | | |
| 3 | Fragmented, double-stranded DNA | 50.00 | 8.33 | 8.33 | |
| 4 | End Repair & A-Tailing Buffer* | 7.00 | 2.33 | 39.67 | |
| 5 | End Repair & A-Tailing Enzyme Mix* | 3.00 | 0.50 | 8.50 | |
| 6 | Total volume: | 60.00 | 20.00 | 2.83 | (per |

4. 30 min 20ºC and 30 min 65ºC.
5. Mix Y adaptor components in a PCR tube. Adaptor sequeces for the different barcodes can be found in Annex Table 1.

**Y adaptor annealing:**

| | A | B | C |
|---|---|---|---|
| 1 | Component | Volume | |
| 2 | ISA## (100µM) | 10.00 | |
| 3 | MiSeq Common Adapter_MI (100µM) | 10.00 | |
| 4 | TE | 80.00 | |
| 5 | Total volume: | 100.00 | |

6. Anneal Y adaptor with the following program: 95°C for 1 s; 60°C for 1s; slow ramp down (approximately -2°C/min) to 4°C; hold at 4°C. Store in -20°C.
7. Mix adaptor ligation reagents in a PCR tube, in the following order:

**Y Adaptor Ligation:**

| | A | B | /3 | /3(x5.5) | E |
|---|---|---|---|---|---|

| | | | | | |
|---|---|---|---|---|---|
| 1 | Water | 5.00 | 1.67 | 9.17 | 4.583333333 3 |
| 2 | DNA Ligation Buffer | 30.00 | 10.00 | 55.00 | 27.5 |
| 3 | DNA Ligation Mix | 10.00 | 3.33 | 18.33 | 9.166666666 7 |
| 4 | ADAPTOR | 5.00 | 1.67 | | #VALUE! |
| 5 | TOTAL | 50.00 | 16.67 | 15.00 | in each |

8. Mix by pippeting several times. Add 50μL of ligation mixture to 60 of end prepped DNA. Incubated 15 min at 20 deg. Purify with 0.7 bead ratio using ampure XP Beads. Elute in 12
9. Set up PCR1 by mixing the following components. Pcr uses phosphorilated primers to protect from lambda exo degradation:

**PCR1**

| | A | B | C | D | E |
|---|---|---|---|---|---|
| 1 | | | | | DNA |
| 2 | | | | | H20 |
| 3 | | | | | ps_P51_F |
| 4 | 94 | 1' | | | ps_#####_ISP_1 |
| 5 | | | | | blocking 10μM (optional,2.5 |
| 6 | 94 | 30" | x30 | | LongAmp Hot Start Taq 2X M Mix |
| 7 | 61 | 30" | | | TOTAL |
| 8 | 65 | 8.30' | | | |
| 9 | 65 | 20' | | | |
| 10 | 4° | hold | | | |
| 11 | | | | | |

10. Add 3μL of lambda exo to 51μL pcR reaction. Digested 3h at 37. 75°C for 10 min
11. used 23μL beads for bead purification (0,75 eth)

**PCR2**

| | A | B | C | D | E |
|---|---|---|---|---|---|
| 1 | | | | | DNA |
| 2 | | | | | H20 |
| 3 | | | | | PBK_P52_F |
| 4 | 94 | 1' | | | PBK_INSERT_GSP2__R |
| 5 | | | | | blocking 10μM (optional,2.5 ul) |

| | | | | | |
|---|---|---|---|---|---|
| 6 | 94 | 30" | x30 | | PRIMERS ONT |
| 7 | 62 | 30" | | | LongAmp Hot Start Taq 2X Master Mix |
| 8 | 65 | 8.30' | | | TOTAL |
| 9 | 65 | 20' | | | |
| 10 | 4º | hold | | | |
| 11 | | | | | |

12. 0,5x beads were used (25μL). Elute in 11μL TE-ph8.
13. Proceed to load sample in the sequencer following PCR Barcoding *Kit* (SQK-PBK$_{004}$).
14. Data analysis can be performed through a web tool or with code deposited in bitbucket and through the web page application INSERTseq.com.

#### Primers/adaptors:

**Table1**

| | A | B |
|---|---|---|
| 1 | ps_P51_F | T*T*T*C*TGTTGGTGCTGATATTGC AATGATACGGCGACCACCGAGATCTACAC |
| 2 | | |

**Annex Table 1: INSERTseq ADAPTORS**

| | A | |
|---|---|---|
| 1 | ISA01 | AATGATACGGCGACCACCGAGATCTACAC AAGAAAGTTGTCGGTGTCTTTGTG TT |
| 2 | ISA02 | AATGATACGGCGACCACCGAGATCTACAC TCGATTCCGTTTGTAGTCGTCTGT TTT |
| 3 | ISA03 | AATGATACGGCGACCACCGAGATCTACAC GAGTCTTGTGTCCCAGTTACCAGG TT |
| 4 | ISA04 | AATGATACGGCGACCACCGAGATCTACAC TTCGGATTCTATCGTGTTTCCCTA TTT |
| 5 | ISA05 | AATGATACGGCGACCACCGAGATCTACAC CTTGTCCAGGGTTTGTGTAACCTT TTT |
| 6 | ISA06 | AATGATACGGCGACCACCGAGATCTACAC TTCTCGCAAAGGCAGAAAGTAGTC TT |
| 7 | ISA07 | AATGATACGGCGACCACCGAGATCTACAC GTGTTACCGTGGGAATGAATCCTT TTT |
| 8 | ISA08 | AATGATACGGCGACCACCGAGATCTACAC TTCAGGGAACAAACCAAGTTACGT TT |
| 9 | ISA09 | AATGATACGGCGACCACCGAGATCTACAC AACTAGGCACAGCGAGTCTTGGTT TT |
| 10 | ISA10 | AATGATACGGCGACCACCGAGATCTACAC AAGCGTTGAAACCTTTGTCCTCTC TT |

15. Tsai, S. Q., Zheng, Z., Nguyen, N. T., Liebers,M., Topkar,V. V., Thapar,V., Wyvekens, N., Khayter, C., Iafrate, A. J., Le,L. P., Aryee,M. J., & Joung, J. K. (2015). GUIDE-seq enables genome-wide profiling of off-target cleavage by CRISPR-Cas nucleases. *Nature Biotechnology*, *33*(2), 187–197.
